# Single-Molecule Tracking of Chromatin-Associated Proteins in the *C. elegans* Gonad

**DOI:** 10.1101/2021.04.04.438402

**Authors:** Lexy von Diezmann, Ofer Rog

**Affiliations:** Center for Cell and Genome Sciences, Salt Lake City UT, USA 84112; School of Biological Sciences University of Utah, Salt Lake City UT, USA 84112

## Abstract

Biomolecules are distributed within cells by molecular-scale diffusion and binding events that are invisible in standard fluorescence microscopy. These molecular search kinetics are key to understanding nuclear signaling and chromosome organization, and can be directly observed by single-molecule tracking microscopy. Here, we report a method to track individual proteins within intact *C. elegans* gonads and apply it to study the molecular dynamics of the axis, a proteinaceous backbone that organizes meiotic chromosomes. Using either fluorescent proteins or enzymatically ligated dyes, we obtain multi-second trajectories with a localization precision of 15-25 nm in nuclei actively undergoing meiosis. Correlation with a reference channel allows for accurate measurement of protein dynamics, compensating for movements of the nuclei and chromatin within the gonad. We find that axis proteins exhibit either static binding to chromatin or free diffusion in the nucleoplasm, and we separately quantify the motion parameters of these distinct populations. Freely diffusing axis proteins selectively explore chromatin-rich regions, suggesting they are circumventing the central phase-separated region of the nucleus. This work demonstrates that single-molecule microscopy can infer nanoscale-resolution dynamics within living tissue, expanding the possible applications of this technique.

## 2. Introduction

The nucleus is the storage compartment for a cell’s genome, but is also dynamically organized to enable many complex processes. These include the duplication, condensation, and partitioning of chromosomes during cell division; the regulation of transcription by chromatin compaction and recruited proteins; and ribosome biogenesis, which occurs in the nucleolus. Meiosis – the specialized cell-division cycle that produces gametes such as sperm, egg and pollen and underlies sexual reproduction – is one such specialized process. During meiotic prophase, parental chromosome pairs are juxtaposed, and undergo DNA breakage and repair to generate physical linkages (crossovers) between chromosomes that are used to correctly partition chromosomes into gametes (Page and Hawley, 2004; Zickler and Kleckner, 1999). Fundamental to meiosis is the unique packaging adopted by meiotic chromosomes, as a series of loops anchored at their base to a protein complex known as the meiotic axis (Fig. 1). This structure is universally conserved, and is essential to almost all aspects of the meiotic program, including the alignment of the parental chromosomes and crossover repair. While extensive analyses have defined the localization of, and binding interfaces between, the protein components of the axis (Kim et al., 2014; Köhler et al., 2017), our understanding of their assembly and dynamic localization during meiosis remains limited.

**Figure 1.**
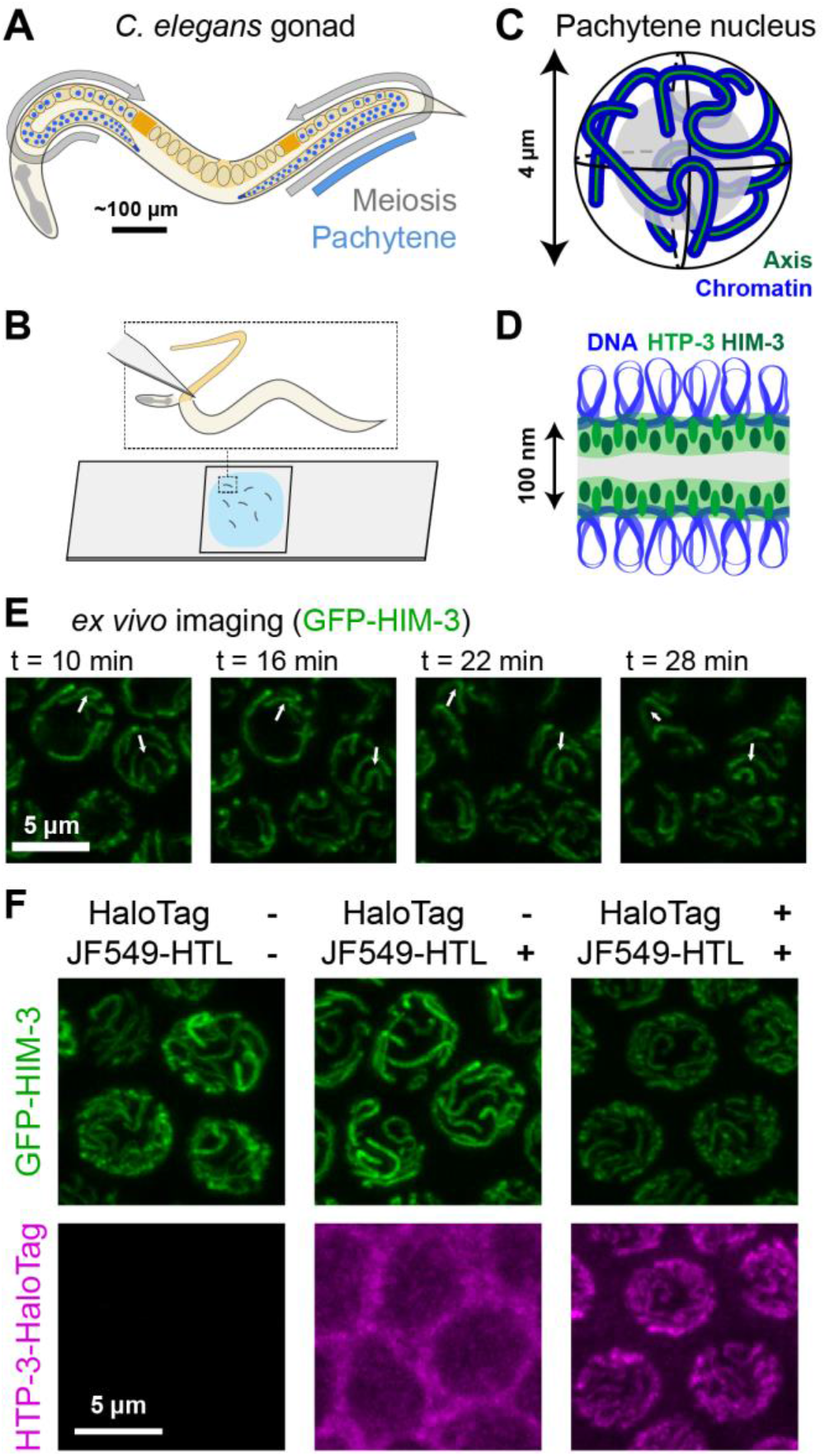
Imaging in the *ex vivo* gonad. A. Schematic of *C. elegans* with both gonad arms marked. Meiosis progresses linearly along the gonad, with the pachytene stage occurring in approximately the middle third. B. Schematic of gonad extrusion. Top, the manual dissection of *C. elegans* causes the gonad to be extruded into solution while remaining supported by attachment to the main body of the worm. Bottom, many extruded gonads are immobilized between an agarose pad and a coverslip mounted on a microscope slide. C. Approximate geometry of the six meiotic chromosomes of *C. elegans* during pachytene. Blue, chromatin; green, axis; gray, nucleolus. D. The nanoscale geometry of looped chromosomal DNA (blue) and the meiotic axis (green). In worms, two major axis protein components are HTP-3 and HIM-3. E. Meiotic nuclei maintain function for tens of minutes following dissection. Images are partial maximum intensity projections of nuclei expressing GFP-labeled HIM-3, which marks the meiotic axis. Arrows highlight the chromosome motion that characterizes this stage of meiosis. F. The gonad is permeable for the synthetic dye JF549 functionalized with a HaloTag ligand (HTL), allowing labeling of HTP-3 fused to the HaloTag enzyme. Partial maximum intensity projections of nuclei are shown for clarity. The image contrast for the GFP and JF549 channels is kept the same between conditions where the HTP-3-HaloTag fusion is present (“HaloTag”) and the dye is or is not added (“JF549-HTL”).

In multicellular organisms, meiosis occurs in specialized organs and tissues, and often in restricted developmental windows. While some of the molecular components and their biochemical activities have been functionally reconstituted *in vitro*, many aspects of their regulation in the context of intact chromosomes remain poorly understood. The nematode *Caenorhabditis elegans* is a particularly convenient system for microscopic studies of meiosis, as it is genetically tractable and the developing gametes within its gonad are ordered in a temporal gradient, allowing different meiotic stages to be easily visualized (Fig. 1A) (Rog and Dernburg, 2013). Moreover, chromosomes in meiotic nuclei are arranged so that they are separate from one another, allowing tracing of individual chromosomes in intact samples. Chemical fixation of extruded gonads for high-resolution microscopy has become a commonplace protocol (Phillips et al., 2009), and super-resolution imaging of such samples has measured the fine structure of the axis and its distinct morphology at different stages of crossover formation (Köhler et al., 2017; Woglar et al., 2020).

In recent years, many advancements have been made by a suite of techniques that probe the dynamics of chromatin-associated proteins (Cho et al., 2018; Chong et al., 2018; Izeddin et al., 2014; Jones et al., 2017; Sabari et al., 2018; Teves et al., 2016). A key component of these efforts has been the application of single-molecule techniques *in vitro* and *in vivo*. Rather than bulk measurements reflecting the average behavior of thousands or more molecules, single-molecule methods reveal underlying heterogeneity that is invisible at the population level (Cuvier and Fierz, 2017). By localizing single molecules in cells, it is possible not only to achieve super-resolution imaging of structure, but also to track the motion of single molecules to detect spatiotemporal dynamics (von Diezmann et al., 2017). While single-particle tracking has predominantly been used in single-celled organisms and cultured cell monolayers, several studies have shown it is possible to extend these studies to *ex vivo* tissue slices (Mashanov et al., 2020; Varela et al., 2016) and developing embryos (Mir et al., 2017; Schaaf et al., 2009). However, these approaches have not been widely applied, possibly due to the need for light-sheet illumination to image within complex tissues (Mir et al., 2017).

In this work, we demonstrate an approach to track single molecules within *C. elegans* gonads, allowing the study of protein dynamics in metazoan meiosis using a standard single-molecule microscope. This development overcomes two major challenges. First, in order to attain high-resolution images while also watching live-cell dynamics, it was necessary to preserve meiotic function in extruded gonad tissue. Second, it was necessary to find labels that were compatible with live-cell imaging and that were sufficiently photostable to provide long tracks, characteristics not guaranteed for many labels used for super-resolution imaging (von Diezmann et al., 2017). Owing to their critical role in meiosis, well-characterized localization and dynamics, and low turnover when bound within a protein complex, we selected the axis proteins HIM-3 and HTP-3 as targets to evaluate our single-molecule tracking approach (Köhler et al., 2017; Rog et al., 2017). Using a nutrient-rich buffer allowed us to preserve meiotic function, while imaging in extruded gonads allowed us to introduce highly photostable dyes optimized for single-molecule imaging (Grimm et al., 2016). We tracked axis proteins with 15-25 nm precision for up to several seconds, and discriminated bound and diffusive modes of motion. By analyzing the subpopulation of freely diffusing molecules, we found that axis proteins are excluded from the nucleolus, suggesting this membrane-less organelle affects the motion of chromatin-associated proteins throughout the nucleus.

## 3. Methods

### Worm strains and transgenes

All strains were cultured using standard methods (Brenner, 1974). Worms were maintained at 20°C except as noted. Strains used were CA1350 (*him-3(ie34[mMaple3::him-3]) IV*) (Rog et al., 2017), ROG118 *him-3*(*gfp::him-3 IV*) (Stauffer et al., 2019), ROG128 (*htp-3(slc2[htp-3::halotag] I)*, ROG213 (*htp-3(slc2[htp-3::halotag]) I*, *him-3(gfp::him-3) IV)*, and CA1330 (*htp-3(tm3655) I; ieSi6 [htp-3::gfp] II; itIs37 [pie-1p::mCherry::H2B::pie-1 3′UTR + unc-119(+)] IV*) (Kim et al., 2015; Wynne et al., 2012). ROG128 (*slc2*, *htp-3::halotag*) was constructed by CRISPR-Cas9 insertion of a *halotag* construct (Los et al., 2008) targeted to the C-terminus of the genomic *htp-3* locus, including a 13 a.a linker (STSGGSGGTGGSS). ROG213 was generated by mating ROG118 with ROG128. As a proxy for the functionality of the tagged axis proteins we quantified hermaphrodite self-progeny (Fig. S1): total progeny and male counts were performed by singling at least 6 healthy L4 hermaphrodites and transferring them to a new plate each day for three days, then counting progeny and males from each plate. HTP-3-HaloTag exhibited meiotic defects, manifesting as lower viable progeny and an increased ratio of males. However, the production of some viable progeny indicates it is at least partly functional. In addition, HTP-3-HaloTag localization was identical to that of untagged HTP-3 (data not shown; (MacQueen et al., 2005)).

### Sample preparation

JF549-PEG-HTL (JF549-polyethylene glycol linker-HaloTag ligand) and PAJF549-HTL (photoactivatable JF549-HaloTag ligand) (gift of Luke Lavis lab, care of Erik Jorgensen lab) were stored at −80°C as 100 μM aliquots in 1 μL DMSO and 20 μM aliquots in 1 μL DMSO, respectively. Immediately before use, aliquots were diluted into M9 minimal medium and then into ECM for labeling during dissections at a final concentration of 10 nM.

Dissection was performed in an adapted embryonic culture medium (Wolke et al., 2007) to preserve native meiotic function. We refer to the medium as ECM, and it is composed of 84% Leibowitz L-15 without phenol red, 9.3% fetal bovine serum, 0.01% levamisole, and 2 mM EGTA, filtered through a 0.2 μm filter and stored in aliquots at −20°C until use. We measured this preparation to have an osmolarity of 330 mOsm (Advanced Instruments Microosmometer 3300), in line with typical osmolarities used when probing the internal environment of *C. elegans* by electrophysiology (Avery et al., 1995). Dissection in these conditions consistently yielded at least a subset of gonads that exhibited meiotic chromosome motion over 10s of minutes. Notably, the original preparation described in (Wolke et al., 2007) includes 4.7% sucrose, which yielded an osmolarity of 500 mOsm and resulted in mild shrinkage of the gonad (data not shown).

To test the ability of minimal medium to support gonad function, we also attempted gonad extrusion in a simple physiological saline buffer without added nutrients (150 mM NaCl, 5 mM KCl, 1 mM MgCl2, 1 mM CaCl2, 10 mM glucose, 15 mM HEPES, pH 7.35, adjusted to 340 mOsm with sucrose). In experiments using this buffer, chromosomes and nuclei did not exhibit persistent motion, indicating meiotic perturbation (data not shown).

For imaging, dissected gonads were placed onto agarose pads. Typically, 2 drops (~30 μL) of 2% melted agarose solution (in water) were dropped to glass slides (VWR Vistavision #16004-422) and flattened by placing another slide on top with thin (~0.2 mm) spacers of lab tape. Agarose pads were stored in a humid chamber at room temperature. To exchange buffer, ECM was added to the pads before use (2 exchanges of 30 μL each, over 10-30 minutes, with excess liquid aspirated between additions and before adding worms).

Worms were dissected following picking of L4 larvae and 16-24 hours’ incubation at 25°C. These staged adult worms were transferred to 20 μL ECM plus 0.01% Tween and, if used, 10 nM dye. Gonads were extruded by incision directly behind the pharynx with Feather #11 scalpel blades. Approximately 10-15 worms were dissected simultaneously and immediately pipetted onto agarose pads, covered with a 22×22 mm^2^No. 1.5 coverslip (VWR #16004-302), and sealed with VaLaP (1:1:1 Vaseline, lanolin, paraffin). Consistent with previous studies (Wolke et al., 2007), gonads that incurred damage during dissection did not exhibit physiological chromosome motions; these gonads were not included in our data analysis.

### Microscopy

Confocal imaging was performed using a Zeiss LSM880 with Airyscan in SR mode, equipped with a 1.4NA 63x oil immersion objective (Zeiss Plan-Apochromat) and using ~49 nm pixel size. Images were acquired with 3-4% 488 nm laser power, 8% 561 nm laser power and 1-2 μs pixel dwell times. All images were Airyscan processed (ZEN Blue 2.3), yielding an approximately 1.7x resolution enhancement over typical confocal resolution.

Single-molecule microscopy was performed on a Bruker Vutara 352 biplane localization microscope (Juette et al., 2008). Briefly: this system provides epi-illumination using collimated diode-pumped solid state lasers sent through a square iris to provide 40×40 μm^2^ square tophat illumination. Data is collected on a Hamamatsu ORCA-Fusion BT sCMOS camera with 99 nm effective pixel size in the image plane. Imaging was performed with a 1.3 NA 60x silicone oil immersion objective (Olympus UPLSAPO60XS2). The irradiances at the image plane used were 0.63 μW/μm^2^ (488 nm), 3.0 μW/μm^2^ (561 nm), and ≤ 0.23 μW/μm^2^(405 nm). PSF calibration, camera read noise calibration, single-molecule biplane localization, and signal photon estimation were performed in the Vutara SRX software using established methods (Huang et al., 2013; Juette et al., 2008).

The radial profile analysis of confocal data was performed using the Radial Profile Extended ImageJ plugin (Philippe Carl, http://questpharma.u-strasbg.fr/html/radial-profile-ext.html).

### Single-molecule data analysis

Calibration of 3D microscope PSFs and single-molecule localization were performed using the Vutara SRX software (Bruker). While the biplane modality of the Vutara 352 estimates the three-dimensional position of single molecules, owing to the relatively poor localization precision in the axial dimension, we analyzed single-molecule data using 2D information only. Track concatenation and mean jump displacement calculations were performed using swift tracking software v. 0.4.3 (Endesfelder, M., Schießl, C., Turkowyd, B., Lechner, T., and Endesfelder, U., manuscript in preparation, software available upon request from https://www.endesfelder-lab.org). This software performs a global fit of single-molecule data, collecting localizations into trajectories in the most likely way for a given set of prior parameters. We used the software parameters: p_bleach = 0.05, exp_noise_rate = 5%, p_blink = 0.1, p_reappear = 0.1, exp_displacement = 150 nm, p_switch = 0.005, precision = 25 nm, diffraction_limit = 150 nm, max_displacement = 700 nm, max_particle_count = 1, and max_blinking_duration = 4 frames. Owing to our choice of activation power and density of labeling, each nucleus generally did not contain more than 1-2 actively fluorescing molecules at a time, making tracks straightforward to discriminate with this approach.

All further analysis was performed in Matlab (The MathWorks, Inc., Natick, MA, USA). Only localizations within nuclei, defined by the diffraction-limited reference channel (Fig. 4B), were used for quantitative analysis.

Mean-squared-displacement analysis was performed as previously described (Lasker et al., 2020). Briefly, we used the 2D diffusion of molecules to estimate diffusivity according to

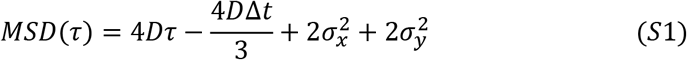

with frame integration time Δ*t* and lag time *τ*, with diffusion coefficient *D* and total localization error 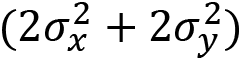 as free parameters determined using nonlinear least-squares estimation. We used tracks with duration of at least 5 frames for this analysis, and performed linear fits to the first four lags of the MSD plot to avoid bias due to higher standard deviations at longer lags (Michalet and Berglund, 2012). In addition to estimates of D described in the text, these fits also yielded estimates of the effective localization precision σ_xy_ of 126 ± 26 nm (mobile molecules, error is standard deviation of bootstrapped tracks) and 15.5 ± 0.5 nm (bound molecules), owing to the different degrees of motion blur in these two populations. While mMaple3-HIM-3 tracks were difficult to analyze in this way owing to their relatively high rate of bleaching, applying similar analysis to tracks with MJDs < 100 nm and duration of 5 or more frames yielded an estimate of σ_xy_ of 27 ± 11 nm. We corroborated this estimate by measuring the empirical localization precision, analyzing the standard deviation of molecule positions in a relatively stationary subset of the trajectory shown in Fig. 2A. In this way, we estimated σ_xy_ to be 24.6 nm (Fig. S3).

**Figure 2.**
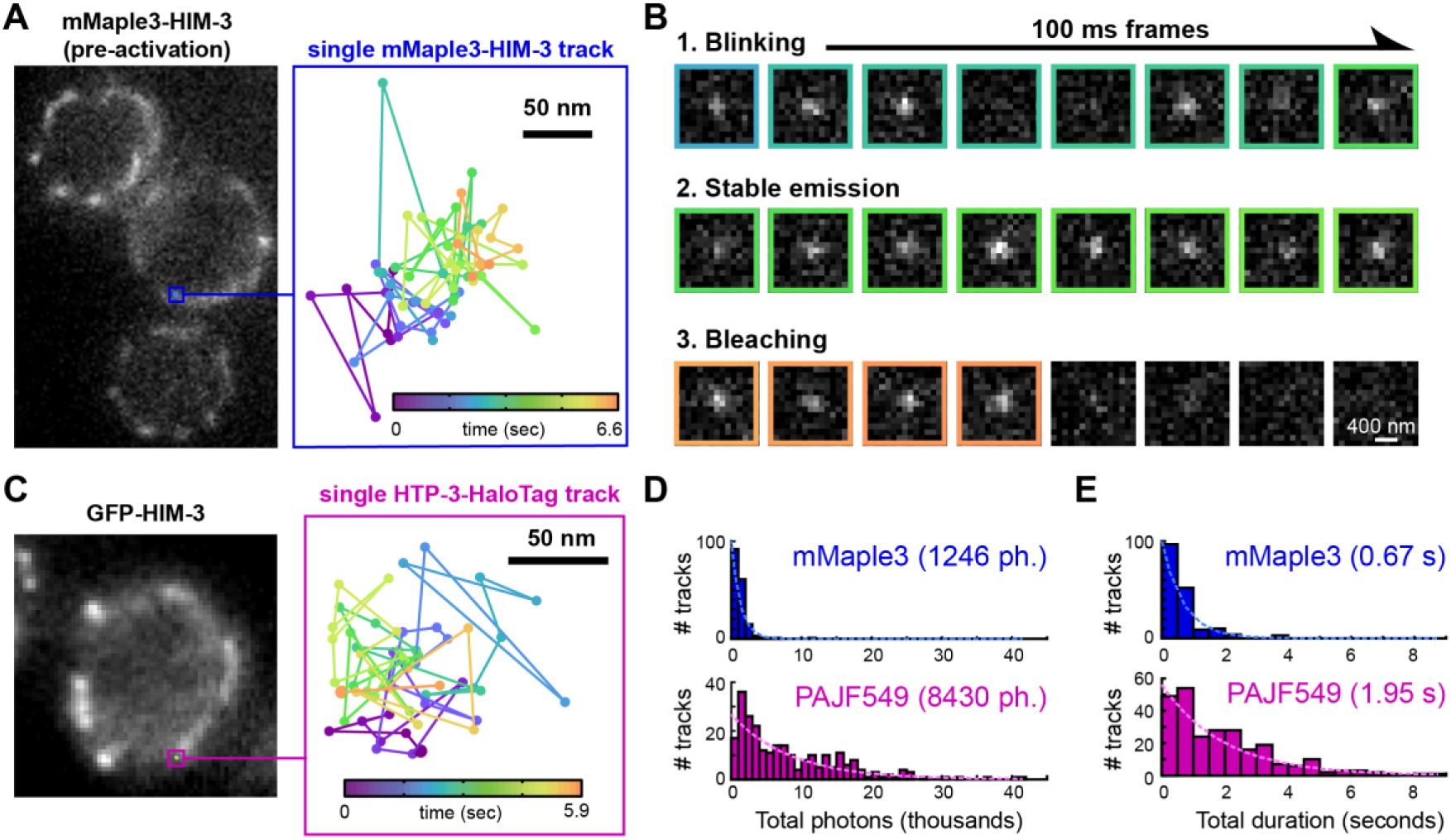
Single-molecule imaging of axis proteins. A. Left, diffraction-limited image of mMaple3-HIM-3 before activation. Right, trajectory of a single photoactivated mMaple-3-HIM-3 molecule associated with the axis. Diffraction limited data are from a single epifluorescence image. B. Selected 100 ms frames of raw data from the molecular trajectory shown in A. Frame borders are colored according to the time colormap in A. One plane of biplane data is shown. C. Left, diffraction-limited image of GFP-HIM-3 reference channel. Right, trajectory of a single photoactivated PAJF549 molecule linked to HTP-3-HaloTag associated with the axis. D. Histogram of total photon counts and corresponding exponential fits for trajectories of mMaple3-HIM-3 and HTP-3-HaloTag (PAJF549), means indicated at top. E. Histogram of total lengths in seconds and corresponding exponential fits for trajectories of mMaple3-HIM-3 and HTP-3-HaloTag (PAJF549), means indicated at top. All imaging performed with 100 ms integration time.

### Registration of diffraction-limited and single-molecule imaging channels

Fluorescent beads (100 nm TetraSpeck™, Invitrogen) were imaged in both the green (reference) and orange (single-molecule) channels. The centers of beads in the reference channel image were mapped to localizations of those beads in the single-molecule channel to generate an affine transformation, which incorporates rotation, scaling, shear, and translation (Fig S4A). The accuracy of this map can be defined by the fiducial registration error (average Euclidean distance between the target positions and positions in the transform); for the data presented here, this value was 23 nm.

## 4. Results

We wished to develop an *ex vivo* imaging protocol that would provide optimal optical properties and dye penetrance, and at the same time maintain physiological meiotic progression (Fig. 1A). Previous imaging of live worms has shown that meiotic chromosomes undergo dynein-driven motion beginning at meiotic entry. Dynein-driven motion invariably ceases upon starvation or when worms are paralyzed with anesthetic (Rog and Dernburg, 2015). Since maintaining a nutrient-rich environment similar to that experienced within animals can preserve oocyte motion in extruded gonads (Fig. 1B) (Wolke et al., 2007), we hypothesized this might also allow meiotic functions to continue. By adapting the embryonic culture medium used in previous work (Wolke et al., 2007), and monitoring GFP-labeled HIM-3 protein to follow the axis (Fig. 1C,D), we found that nuclei exhibited motion within the gonad and that chromosomes dynamically rearranged for up to an hour after dissection (Fig. 1E and Movie S1). Meiotic functions depended on the presence of nutrients available in the culture medium, as nuclei in gonads extruded in physiological saline halted chromosome motion within a few minutes (data not shown). This suggests that overall gonad physiology, and thus meiotic function, was preserved under these conditions.

In addition to lowering autofluorescence and optical aberrations, imaging in live extruded gonads exposes the tissue of the gonad to the surrounding buffer. This is attractive for bioorthogonal labeling approaches such as the HaloTag system, in which a genetically encoded enzyme (the HaloTag enzyme) covalently binds the chloroalkyl moiety (HaloTag ligand, HTL) of an added dye (Los et al., 2008). We tested whether we could introduce dyes into the gonad without extending the time required for sample preparation using the dye JF549-HTL (Lavis et al., 2016) to label endogenously expressed HTP-3-HaloTag. Fully replacing native HTP-3 with the HTP-3-HaloTag partially perturbed meiotic function, but did not disrupt the localization of HTP-3 (Methods, Fig. S1). While the addition of dye to the dissection buffer led to background fluorescence when the HaloTag enzyme was not present, expression of endogenous HTP-3-HaloTag fusion protein effectively quenched this excess dye without requiring a washout step or other extension of our protocol. Notably, while similar bioorthogonal imaging approaches have been successfully used for super-resolution imaging in whole worms, these have required the use of >1000-fold higher dye concentrations and wash-out steps lasting several hours (Li et al., 2016b). This robust and versatile labeling approach expands the toolkit for single-molecule imaging, joining fluorescent tags previously used in worms such as the green-to-red UV-photoconvertible protein mMaple3 (Köhler et al., 2017; Rog et al., 2017; Wang et al., 2014).

We characterized the suitability of photoconvertible mMaple3 and the UV-photoactivatable (dark-to-bright) PAJF549 dye (Grimm et al., 2016) for single-molecule imaging in live gonads. We first tested the blinking properties and photostability of mMaple3 fused to HIM-3. By correlating trajectories of photoconverted single molecules with diffraction-limited images of bulk unconverted mMaple3, we confirmed that detected single molecules appeared primarily within the axis (Fig. 2A). Examining individual trajectories that appeared within the axis complex, mMaple3-HIM-3 was detected for up to several seconds (Fig. 2A) with a localization precision σ_xy_ of ~25 nm (Methods, Fig. S3). These trajectories exhibited digital photoactivation and photobleaching events, as well as occasional brief (≤ 400 ms) blinking events (Fig. 2B, Fig. S2A,B). Consistent with previous work using confocal microscopy (Rog and Dernburg, 2015), the visible laser light we used for single-molecule imaging did not appear to impair meiotic activity, as nuclei and chromosomes continued to exhibit motion during multi-minute acquisitions (Methods, Fig. 2C and Movie S2). Despite this motion, sampling chromosome motion at 30 second intervals was sufficient to contextualize track positions, consistent with previously described root mean squared displacements of ≤ 1.0 μm/minute for chromosomes during the pachytene stage of meiosis (Fig. S4B) (Wynne et al., 2012).

We characterized the photophysics of the PAJF549-HTL dye bound to HTP-3-HaloTag using GFP-HIM-3 as a diffraction-limited reference channel (Fig. 2C). Crucially, PAJF549 afforded dramatically longer tracks with a greater total number of photons (a mean of 2 seconds with 100 ms integration time and 8,400 photons) compared with mMaple3 (0.7 seconds and 1,200 photons) (Fig. 2D,E, Fig. S2C). Additionally, PAJF549 could be consistently localized for a larger fraction of each track: for mMaple3, 43% frames of each track were marked as too dim to localize due to intensity fluctuations (blinking), while for PAJF549, only 22% of frames per track were not possible to localize accurately (Fig. S2D). These results are consistent with the greater photostability of photoactivatable dyes relative to fluorescent proteins *in vitro* and in other biological systems (Lavis et al., 2016; Lee et al., 2010; Wang et al., 2014).

The longer single-molecule trajectories that we obtained for PAJF549-labeled HTP-3-HaloTag molecules allowed us to discriminate between several subpopulations of molecules. Overlaying HTP-3-HaloTag trajectories with the reference channel (Fig. 3A) suggested two such populations: a large fraction of static molecules that co-localized with the axes, and freely diffusing molecules that did not exhibit correlation with GFP-HIM-3. To quantify these populations, we calculated the mean jump displacements of each molecule trajectory and analyzed the resulting distribution (Fig. 3B). This distribution exhibited an inflection point at a mean jump displacement of ~75 nm. We interpret this as a bound fraction of HTP-3 molecules integrated into the axis (two-thirds of all detected molecules) and a mobile fraction of free molecules (one-third). Since no single cut-off perfectly separated this bimodal distribution, we defined windows for tracks with mean jump displacements of < 50 nm and of > 100 nm as bound and mobile, respectively. By separately applying mean squared displacement (MSD) analysis to each subpopulation (Fig. 3C), we found that bound molecules exhibited only small amounts of confined motion (D = 5×10^−4^± 1×10^−4^μm^2^/s at short times, and flattening MSD curves at longer lag times). We estimated a value for the PAJF549 localization precision of σ_xy_ = 15.5 ± 0.7 nm from the MSD curve (Methods). The mobile fraction exhibited Brownian diffusion with diffusion coefficient D = 0.10 ± 0.04 μm^2^/s, exemplified by the linear MSD curve even at longer intervals (Fig. 3C). These measurements are in the range of diffusion coefficients measured by single-molecule tracking of transcription factors and slower (as expected) than those measured for fluorescent proteins not fused to nuclear proteins (Izeddin et al., 2014; Li et al., 2016a). These results were only weakly sensitive to the choice of mean jump displacement cutoff used to categorize subpopulations, with values of D increasing 4% for the bound population when the lower threshold was raised to 75 nm, and decreasing 21% for the mobile population when the upper threshold was lowered to 75 nm.

**Figure 3.**
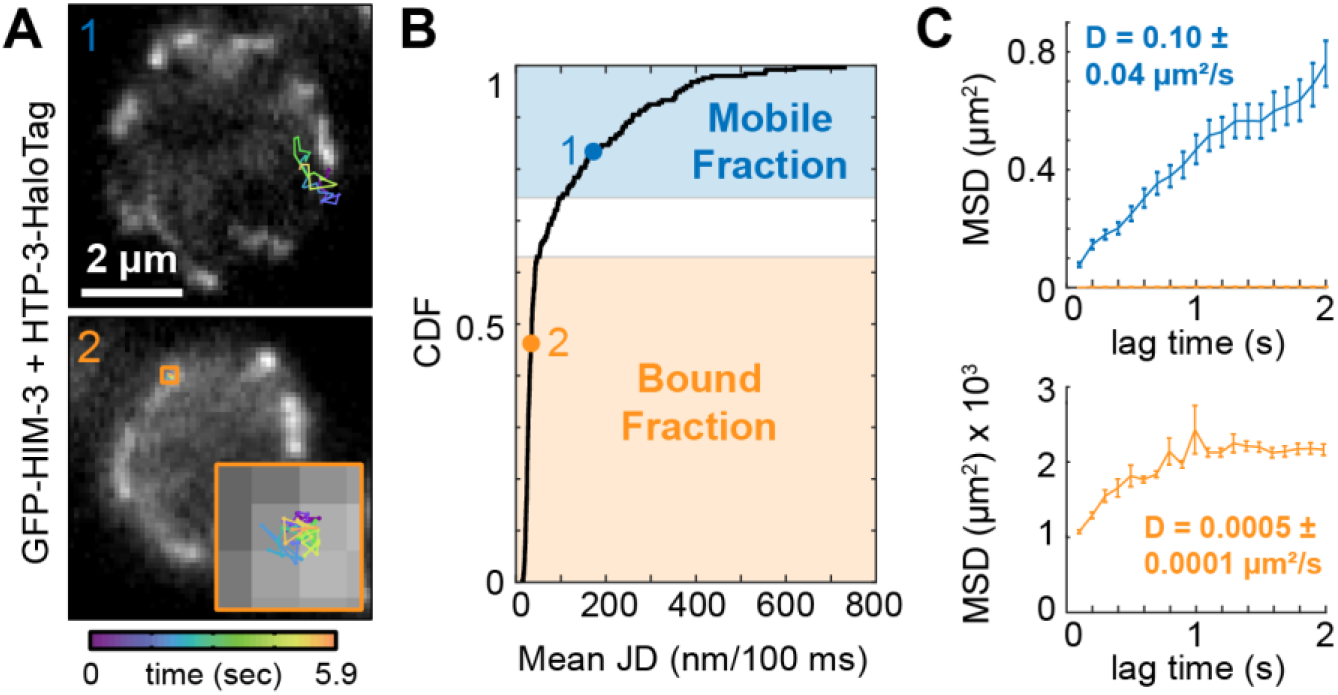
Single-molecule tracking of HTP-3 subpopulations with different dynamics. A. Single-molecule trajectories of HTP-3-HaloTag molecules overlaid onto diffraction-limited reference images of GFP-HIM-3, marking the axis in pachytene nuclei. Top, a trajectory displaying diffusion throughout the nucleoplasm. Bottom, a static trajectory within a single axis. B. the cumulative distribution function (CDF) of mean jump displacements (JDs) for HTP-3-HaloTag molecules. Dots on the CDF curve mark the mean jump displacements of the two tracks shown in A. C., mean squared displacements as a function of lag time for the mobile and bound trajectory populations marked in B. Top, mobile fraction. The bound fraction is also plotted for reference, appearing virtually identical to the x axis at this scale. Bottom, the bound fraction, with MSD scaled by a factor of 1000. Error bars of each MSD value represent standard error. Precision of D estimate is the standard deviation when bootstrapping tracks.

Our observation of Brownian diffusion for mobile HTP-3-HaloTag molecules suggests motion was largely unobstructed, at least on the ≤ 1 μm scale (Fig. 3C). On larger scales, however, diffusion is likely to be affected by the heterogeneity of the nucleoplasm, which includes, among many other characterized compartments, chromosomes, heterochromatin, and the nucleolus. The latter is perhaps best characterized for its ability to selectively recruit client macromolecules while excluding condensed chromatin and other macromolecules (Lafontaine et al., 2020).

To investigate whether HTP-3 molecules were able to freely explore the entire nucleus, we first examined bulk localization in premeiotic nuclei, where HTP-3 is expressed but has not yet associated with chromatin to form the meiotic axis. Confocal slices of nuclei immunolabeled for HTP-3 alongside DAPI and the nucleolar marker DAO-5 (Lee et al., 2014) allowed us to demarcate the localization pattern of HTP-3 relative to chromatin and to the nucleolus, which in the worm germline forms a sphere filling the middle of the nucleus (Fig. 1C, Fig. 4A, top). Averaging protein distributions from multiple nuclei within a shared coordinate system did not show a significant difference of HTP-3 density in the nucleolus (Fig. 4A, top). However, as has been shown for transcription factors in embryonic stem cells, fixation can bias the observed distribution of chromatin-associated proteins away from DNA (Teves et al., 2016). Repeating this experiment in live gonads using genetically encoded mChy-H2B at a marker for chromatin, we found that GFP-HTP-3 exhibited clear depletion from the center of the nucleus (Fig. 4A, bottom), indicating that prior to axis assembly, axis proteins are partitioned away from the nucleolus and into chromatin-rich regions.

**Figure 4.**
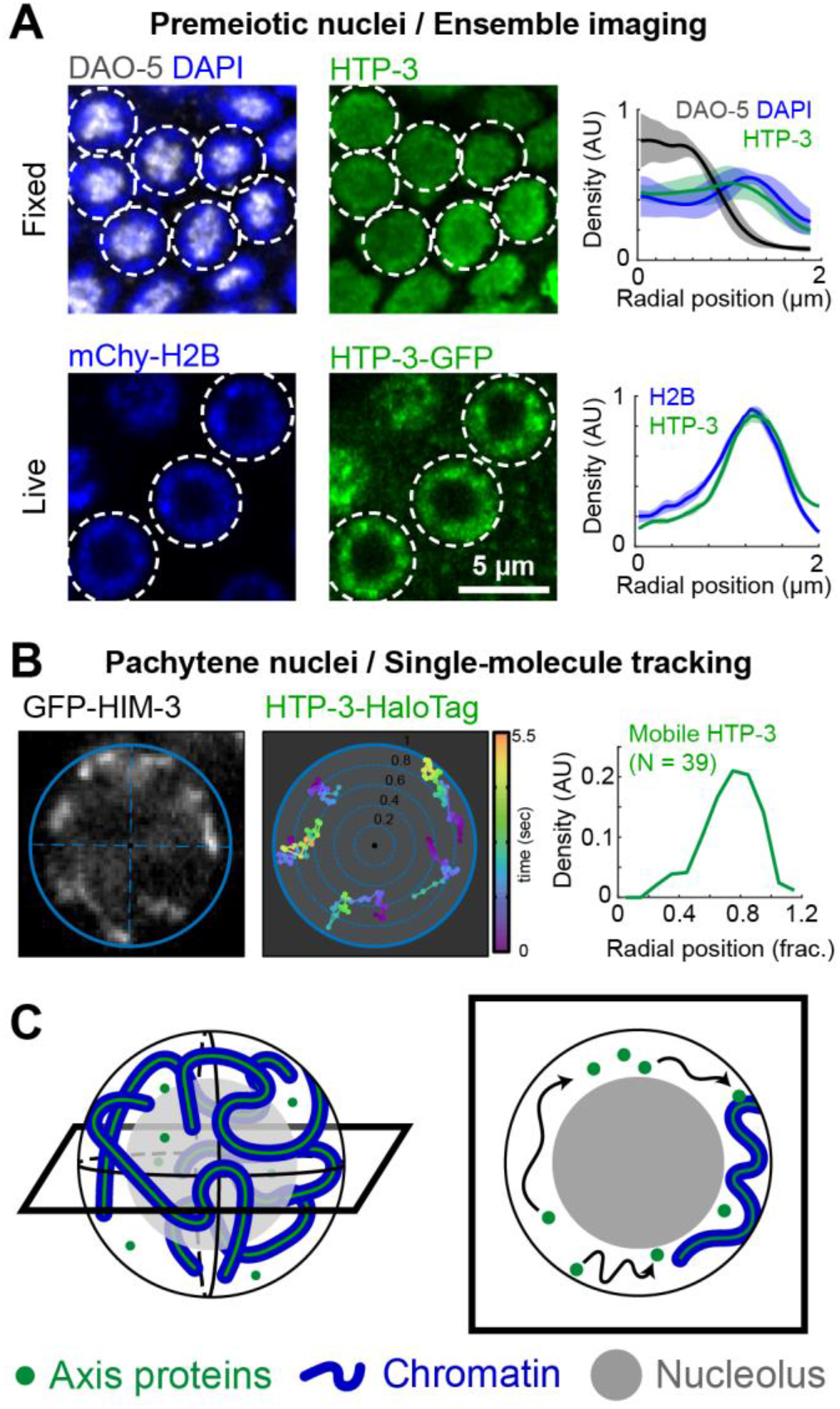
The nucleolus excludes HTP-3 molecules. A. Confocal imaging of free axis proteins (HTP-3) relative to DNA (DAPI, mChy-H2B) and the nucleolus (DAO-5) in premeiotic nuclei. Top, immunolabeling of fixed gonads. Bottom, imaging in live extruded gonads. White circles: nuclei selected to generate profiles of radial density, shown on the right. Each line, mean of density profiles in indicated nuclei; shaded areas, standard deviation. B. Distribution of single-molecule tracks of HTP-3 in meiotic (pachytene) nuclei. Left, example ellipse drawn onto diffraction-limited GFP-HIM-3 reference image to define nuclear coordinates. Middle, five tracks registered within the shared polar coordinate system. Right, distribution of mobile HTP-3 molecule localizations from 39 tracks. C. Schematic of axis protein dynamics within pachytene chromosomes. Left, three-dimensional nuclear geometry, with cross-section shown at right. Mobile axis proteins are excluded from the nucleolus and must transit around the nuclear periphery (black arrows) to find binding sites.

Our single-molecule approach allowed us study a more physiologically relevant scenario: the diffusion of unbound molecules at a time when a protein of interest exhibits a well-defined localization pattern on chromatin. We returned to pachytene nuclei and used our single-molecule data to examine the specific case of HTP-3 molecules that were not associated with chromatin (the mobile fraction as defined in Fig. 3). We found that these tracks were preferentially partitioned away from the nucleolus, with a distribution similar to that in the premeiotic, live-cell case (Fig 4B). This suggests that chromatin-associated proteins are selectively excluded from the nucleolus, requiring their diffusion to circumvent the center of the nucleus and specifically traverse chromatin-rich regions. Nucleolar occlusion could impact search kinetics: in the case of *C. elegans* nuclei, search would occur in a shell at the periphery of the nucleus rather than throughout the full nuclear volume (Fig. 4C).

## 5. Discussion

In this work, we established an approach to image extruded *C. elegans* gonads. This system allowed us to probe nuclei actively undergoing meiosis using single-molecule imaging. In addition to avoiding autofluorescence common when imaging whole animals, our method allowed us to use a variety of newly-developed labels and synthetic dyes. We quantitated the photophysics of both mMaple3 and HaloTag-bound PAJF549 when fused to proteins of the meiotic axis, and found that while both labels were suitable for single-molecule tracking, the greater photostability of PAJF549 supported categorization of heterogeneous molecular diffusivities. Sifting tracks by diffusivity allowed us to analyze the distribution of mobile, free molecules of the axis protein HTP-3, and to show that their diffusion trajectories sidestep the nucleolus.

Successful execution of the meiotic program requires dramatic remodeling of chromosomes and tightly-regulated production of chromosomal crossovers, as described in e.g. (Conrad et al., 2008; Koszul et al., 2008; Marshall et al., 2016; Rog and Dernburg, 2015; Wynne et al., 2012). While these dynamic processes are integral to meiosis, super-resolution imaging of chromosome structure and DNA repair intermediates has been limited to fixed samples (Köhler et al., 2017; Woglar and Villeneuve, 2018), and there have been only a few studies investigating molecular turnover within meiotic structures (Stauffer et al., 2019). In this work, we apply single-molecule tracking to study the dynamic behavior of chromatin-associated molecules. This revealed the underlying flow of a population of freely diffusing molecules that was previously invisible. The *ex vivo* gonad preparation we have optimized here allowed us to use newly-developed dyes and an optically advantageous sample while maintaining physiological meiotic function. Future work, taking advantage of the transparency and size of *C. elegans*, recent progress in immobilization procedures (Burnett et al., 2018; Rog and Dernburg, 2015) and the development of novel synthetic dyes and florescent proteins (Li and Vaughan, 2018), should enable to extend our approach to live, intact animals.

Single-molecule tracking has shown that chromatin-associated proteins such as transcription factors exhibit multiple search types (e.g., one-dimensional diffusion along chromatin, transient interactions, and three-dimensional diffusion), and that these are affected by the local nuclear environment (Brown et al., 2016; Izeddin et al., 2014; Li et al., 2016a). Our approach promises to expand the scope of these studies to complex biological processes that occur in whole tissues and animals. Some of the meiosis-specific processes that are likely to benefit from the rich information generated by single-molecule tracking include chromatin remodeling during axis assembly (Kim et al., 2014), the liquid-like organization of the synaptonemal complex (Nadarajan et al., 2017; Pattabiraman et al., 2017; Rog et al., 2017), and the recruitment of repair proteins to prospective crossover sites (Zhang et al., 2018).

The concept of biomolecular phase separation has recently emerged as a provocative new framework to understand cellular organization and signaling. Phase separation in the nucleus appears to play a role in diverse aspects of biology including development, neuronal function, and immunology (Alberti, 2017; Chen et al., 2020; Yoo et al., 2019). Indeed, fundamental components of the nucleus can be viewed as phase-separated entities, and recent work has shown that a diffusion boundary exists between euchromatin and heterochromatin (Strom et al., 2017) and that the nucleolus is composed of multiple phase-separated subdomains (Lafontaine et al., 2020). However, with few exceptions, these studies have focused on the composition of phase-separated membrane-less organelles and the consequences of sequestration within them. Here we have shown that membrane-less organelles also exert control over their surroundings: by being unable to enter the nucleolus, chromatin-associated proteins must traverse the nuclear periphery, effectively sequestering and enriching them within chromatin-rich regions. The application of single-molecule tracking promises to reveal further implications of the multifarious phase-separated entities within cells.

## 6. Supplementary Information

**Figure S1.**
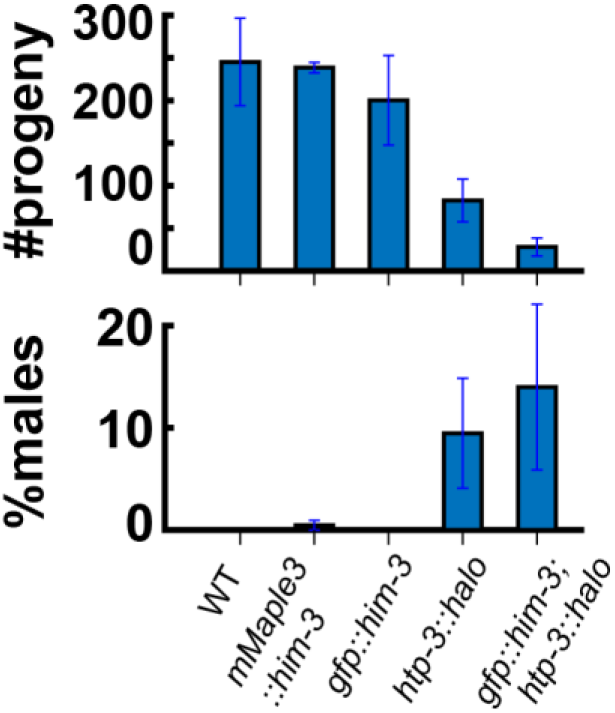
Progeny counts of strains used in this study. Reductions in progeny reflect defects in meiosis, and an increase in males in the population specifically reflects X chromosome nondisjunction. Error bars reflect a 95% confidence interval (N ≥ 5hermaphrodites for each condition).

**Figure S2.**
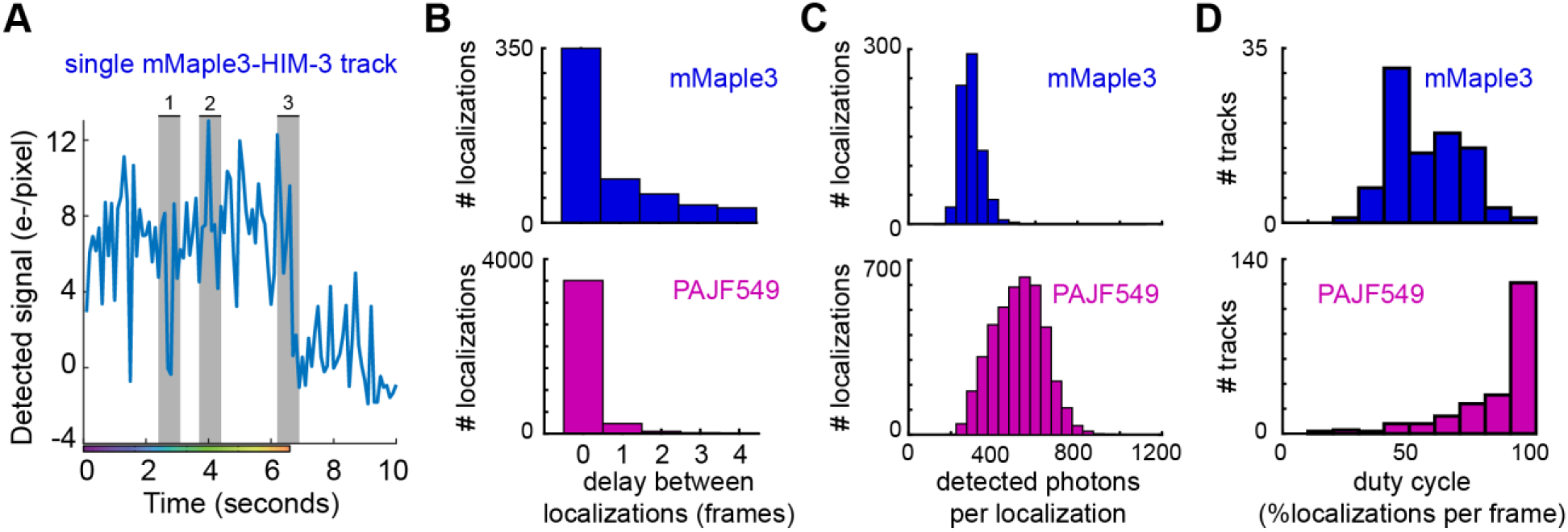
Photophysical characterization of mMaple3 and PAJF549. A. raw intensity data (signal above background) for the single-molecule mMaple3-HIM-3 track shown in Fig. 2. The colormap shown corresponds to the segment of the track displayed in Fig. 2A-B. The sections marked 1, 2, and 3 correspond to the three parts of the track referred to ‘blinking’, ‘stable emission’ and ‘bleaching’. B. the delay between successful localizations for mMaple3 and PAJF549. C. the photon counts per localization (i.e., each frame) for mMaple3 and PAJF549. D. the duty cycle (measured as the percentage of detected localizations per frame of a trajectory) for tracks of duration 5 or more frames.

**Figure S3.**
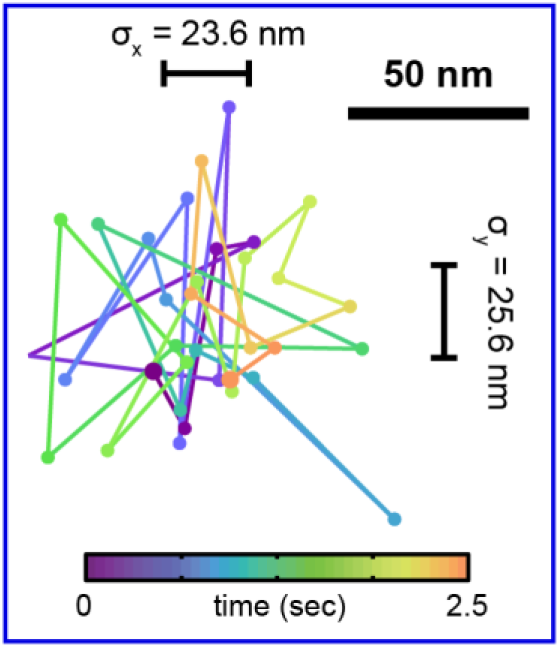
A subset of the track displayed in Fig. 2A that does not display overall motion (enlarged dots: beginning and end of trajectory). The standard deviation of localizations in this subset is 23.6 nm in *x*, 25.6 nm in *y*, for an estimate of σ_xy_ of 24.6 nm (geometric mean).

**Figure S4.**
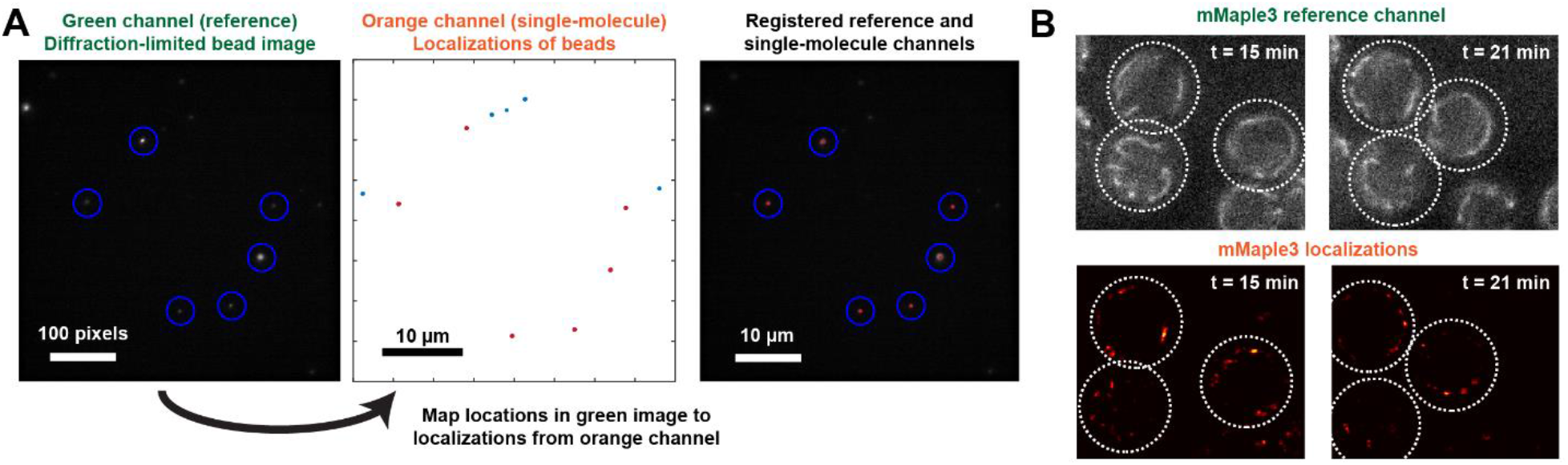
Registration of single-molecule and diffraction-limited channels. A. To use the green diffraction-limited channel as an accurate reference for single-molecule data collected in the orange channel, we generate an affine transformation matrix between image space and localization space using multispectral beads (Tetraspeck, 100 nm). We select the centers of the bead images in the green channel (blue circles) and the average position of repeatedly localized beads in the orange channel (red dots; blue dots, localizations not selected) to generate this transformation. Applying this transformation to the reference channel places it in the same coordinate system as the single-molecule data (fiducial registration error for this dataset: 23 nm). B. Sampling the reference channel at frequency 1/minute allows us to contextualize track positions in the nucleus. Top, mMaple3-HIM-3 reference images at 15 minutes and 21 minutes after dissection. Bottom, reconstructed images of all mMaple3-HIM-3 single-molecule localizations acquired in 1-minute windows centered at 15 minutes and 21 minutes after dissection. Note movement of the nuclei relative to each other, as well as rearrangements of the chromosomes inside the nuclei.

**Movie S1.**
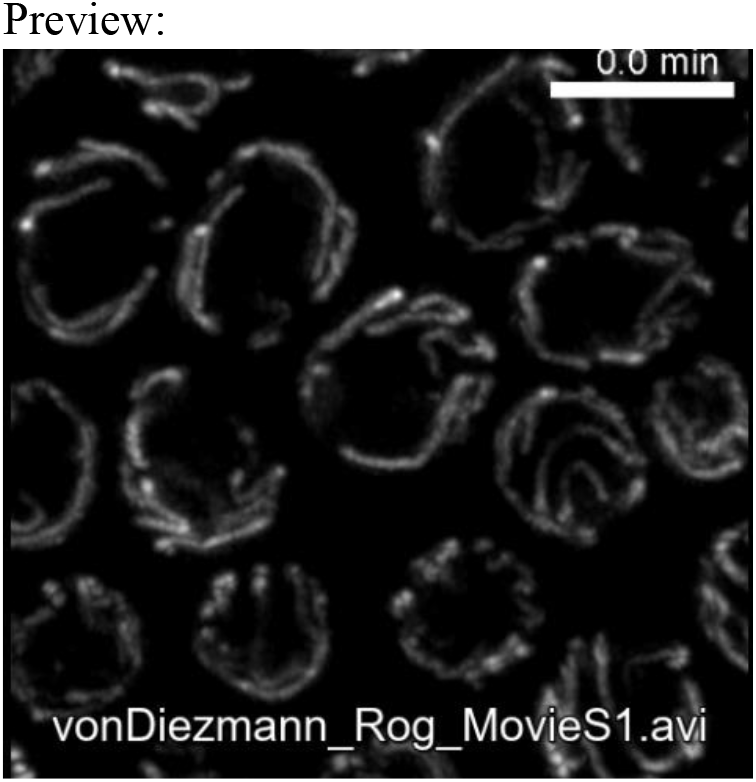
Confocal images of GFP-HIM-3 taken at 3 minute intervals. Scalebar, 5 μm.

**Movie S2.**
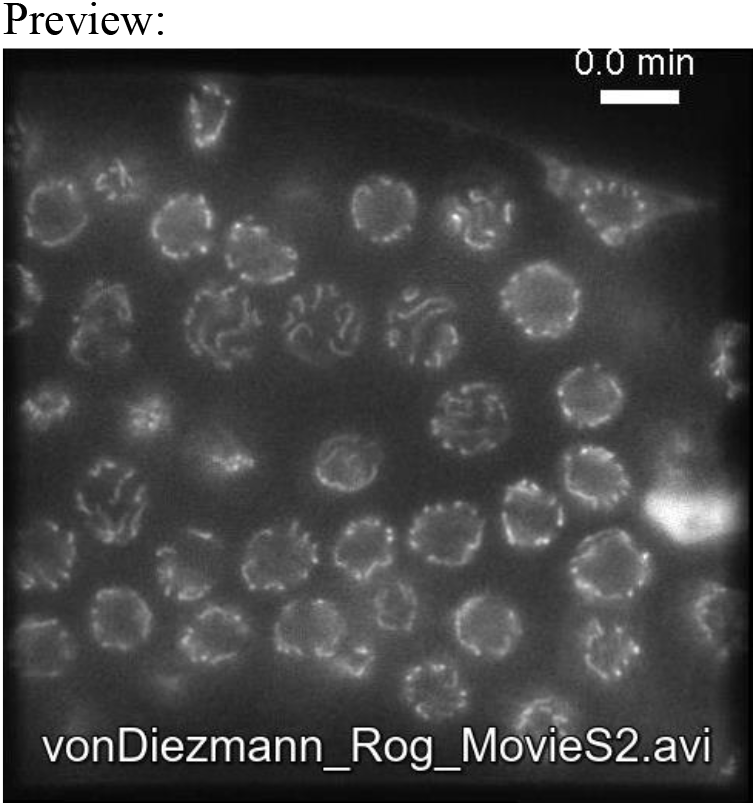
Widefield epifluorescence images of GFP-HIM-3 taken at 30 second intervals, interleaved with single-molecule imaging of HTP-3-HaloTag (not shown). Scalebar, 5 μm.

## 7. Acknowledgements

We thank Abby Dernburg for worm strains, the Rog and Erik Jorgensen labs for discussions, and Sara Nakielny for comments on the manuscript and editorial work. We thank Presley Azarcon for contributions to worm strain generation early in this project. We thank Thien Vu and Siyu Chen for assistance with reagents and osmometry. We are grateful to Jonathan Grimm and Luke Lavis for the generous gift of fluorescent dye samples. We thank the Jorgensen lab and Bruker (Rob Hobson) for the use of, and assistance with, the Vutara 352 single-molecule microscope. L.v.D. is The Mark Foundation for Cancer Research Fellow of the Damon Runyon Cancer Research Foundation (DRG-2372-19). This work was supported in part by funding to O.R. from the University of Utah and from NIGMS (NIH) under award number R35GM128804.

